# Directing an mRNA-LNP vaccine toward lymph nodes improves humoral and cellular immunity against SARS-CoV-2

**DOI:** 10.1101/2021.08.25.457699

**Authors:** David M. Francis, Runqiang Chen, Sahba Khorsandzadeh, Qidong Hu, Xiaoxuan Lyu, Hua Wang, Wan-lin Lim, Haotian Sun, Hui Xie, Namir Shaabani, Russell Ross, Brian Cooley, Henry Ji

## Abstract

The exploration and identification of safe and effective vaccines for the SARS-CoV-2 pandemic has captured the world’s attention and remains an ongoing issue in order to protect against emerging variants of concern (VoCs) while generating long lasting immunity. Here, we report the synthesis of a novel messenger ribonucleic acid (mRNA) encoding the spike protein in a lipid nanoparticle formulation (LNP) (STI-7264) that generates robust humoral and cellular immunity following immunization of C57Bl6 mice. In efforts to continually improve immunity, a lymphatic drug delivery device (MuVaxx) was engineered and tested to modulate immune cells at the injection site (epidermis and dermis) and draining lymph node (LN) to elicit adaptive immunity. Using MuVaxx, immune responses were elicited and maintained at a 10-fold dose reduction compared to traditional intramuscular (IM) administration as measured by anti-spike antibodies, cytokine producing CD8 T cells, and neutralizing antibodies against the Washington (Wild Type, WT) and South African (beta) variants. Remarkably, a 4-fold elevated T cell response was observed in MuVaxx administered vaccination as compared to that of IM administered vaccination. Thus, these data support further investigation into STI-7264 and lymphatic mediated delivery using MuVaxx for SARS-CoV-2 and VoCs vaccines.

## Introduction

The SARS-CoV-2 virus has accounted for more than 210 million cases of the coronavirus disease 2019 (COVID-19) and over 4.4 million fatalities worldwide since its original outbreak in December 2019. This is the 3^rd^ outbreak of a *Betacoronavirus* since 2002, the SARS-CoV and Middle East respiratory syndrome coronavirus (MERS-CoV) being its predecessors, and is much more efficiently transmitted person to person. While vaccine efforts have greatly hampered the spread of disease, questions about their durability and protection against emerging VoCs remain.^1,2^ Thus, there is an urgent need to improve vaccine durability and efficacy to emerging VoCs, while balancing costs, stability, and manufacturing speed to scale up for world-wide vaccination efforts.

There are currently two mRNA-based SARS-CoV-2 vaccines which have been authorized and widely disseminated. These vaccines encode for the spike (S) protein, which is the major surface protein on the coronavirus virion and are thus the primary target for neutralizing antibodies (nAb) as the S protein facilitates viral entry into host cells via interactions with the angiotensin-converting enzyme 2 (ACE2) receptor expressed in the upper and lower respiratory tract. The primary metric for SARS-CoV-2 vaccine efficacy has been on generating nAbs to prevent viral entry into host cells and promote viral clearance before replication. In addition to humoral immunity, an ideal vaccine would also generate cellular immunity as T cell immunity has been associated with less severe disease,^3,4^ faster recovery,^4^ and memory persistence for decades.^5^

To date, most vaccines are administered IM due to feasibility for healthcare workers, speed of injection, and immunological properties (i.e. muscle resident lymphocytes and antigen presenting cells). However, directing vaccines toward the dermis and draining LNs has been of interest for decades due to the high concentration of antigen presenting cells (APCs) including Langerhan cells that reside in the skin (epidermis and dermis) that are capable of taking up antigen and subsequently trafficking to draining LNs to elicit adaptive immunity.^6,7^ Moreover, the initial lymphatics are present at high concentrations just below the stratum corneum and provide direct access to draining LNs due to their high permeability and uni-directional flow towards draining LNs.^8,9^ Delivering vaccines directly to LNs provides a promising opportunity for improving vaccine efficacy as 1) lymphocytes reside in LNs at high concentrations,^10^ 2) are home to unique and strategically positions APCs that present incoming antigen to T and B cells rapidly to induce immunity,^11,12^ and 3) memory T and B cells reside in LNs during their lifespans.^13,14^

To address the SARS-CoV-2 pandemic and need for effective vaccines, we engineered a novel mRNA construct encoding the S protein in a lipid nanoparticle formulation (LNP) referred to as STI-7264 and interrogated the immunological response using a traditional intramuscular (IM) injection compared to our proprietary Sofusa MuVaxx Lymphatic Drug Delivery Platform (MuVaxx). We first explored the ability of MuVaxx to deliver antigen to LNs to induce an improved immunological response by vaccinating against a model antigen, Ovalbumin (OVA) in a preclinical mouse model compared to an IM injection. We next compared STI-7264 against a marketed reference mRNA-LNP (Reference) vaccine for comparison and characterized the humoral and cellular response in a preclinical mouse model measuring circulating anti-S antibodies and peripheral cytokine producing T cells. Our STI-7264 formulation led to similar antibody production compared to the Reference vaccine, however the CD8 T cell response was dramatically improved. MuVaxx administration of STI-7264 enabled an approximately 10-fold reduction in dose needed compared to an IM injection while maintaining similar B cell immunity and an elevated T cell immunity. Thus, STI-7264 and MuVaxx represent an intriguing approach for vaccinating against SARS-CoV-2 and its VoCs as well as moving forward to clinical applications.

## Results

### STI-7264-SARS-Cov-2 mRNA is an mRNA vaccine for prevention of COVID-19

The vaccine is comprised of an active drug substance, a single-stranded mRNA encoding for the full-length SARS-CoV-2 S glycoprotein encapsulated in lipid nanoparticles (LNPs). The sequence was derived from the strain “Severe acute respiratory syndrome coronavirus 2 isolate Wuhan-Hu-1”. Mutations were introduced into S protein to substitute residues 986 and 987 to produce prefusion-stabilized SARS-CoV-2 S(2P) protein.^15^ To achieve optimal expression in humans, the sequence was further codon-optimized and cloned into a pVAX1-based backbone that contains T7 promoter, 5′-UTR, 3′-UTR and optimized Poly-A tail with minimal overhang. The template was then linearized immediately downstream of the Poly-A tail and used for in vitro transcription (IVT) (Figure 1A). To facilitate mRNA expression and reduce innate immune response, during the IVT, Cap 1 structure was added to the 5’ terminus of the RNA co-transcriptionally by CleanCap® AG, and UTP completely replaced by N1-methylpseudo-UTP. This process can be readily scaled up to produce desired amounts of capped mRNA.

**Figure 1).**
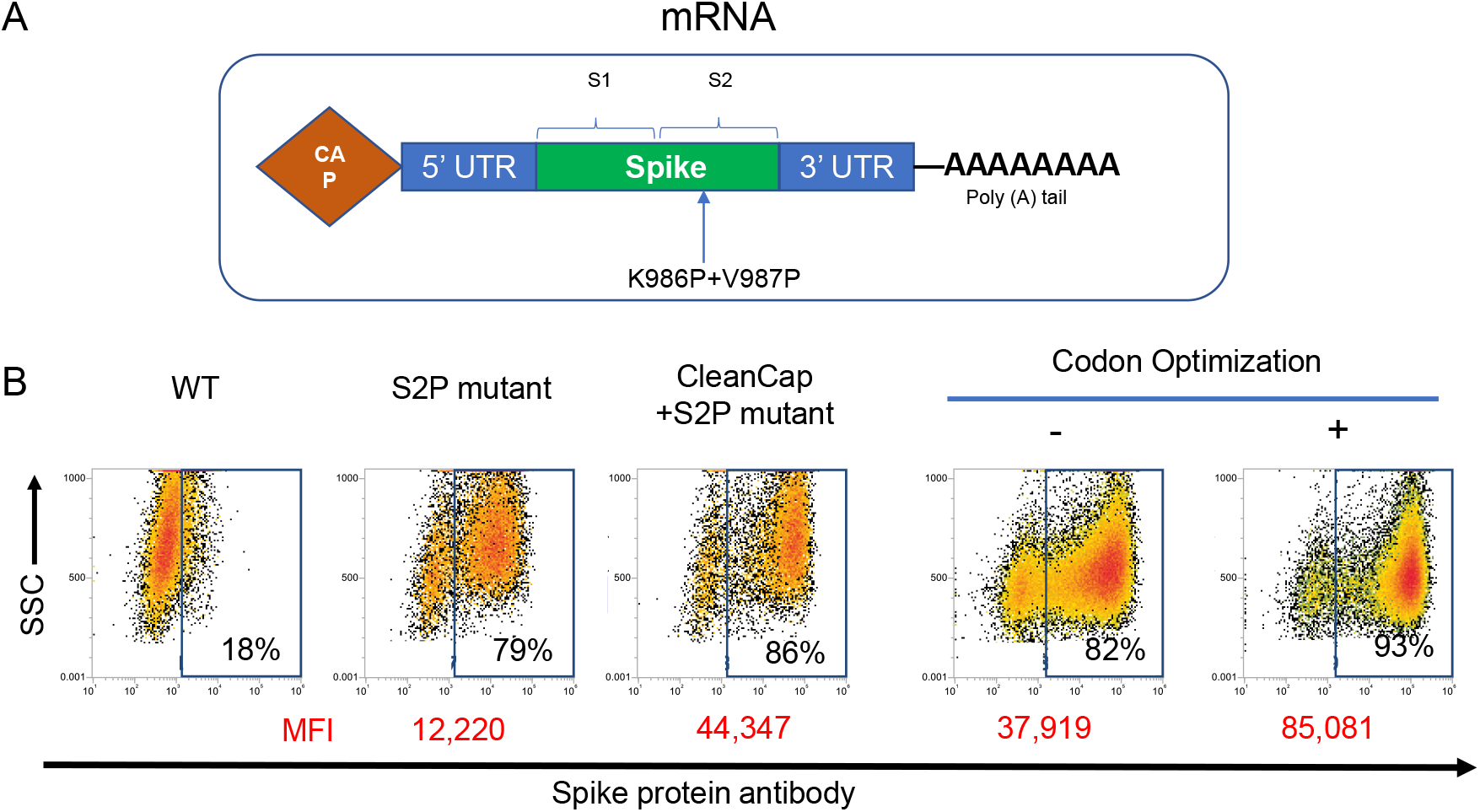
STI mRNA vaccine is optimized for highly efficient translation. A) Schematic for STI mRNA sequence. B) Primary dendritic cells were transfected with various mRNA, stained with anti-Spike antibody STI-2020, and evaluated by flow cytometry 24 post-transfection.

To confirm the expression, the IVT mRNA was introduced into monocyte-derived dendritic cells (DC) by electroporation. Twenty-four hours post-transfection, the cells were collected and stained with anti-Spike antibody STI-2020 and detected with allophycocyanin-conjugated anti-human Fc antibody. The flow cytometry showed that the codon-optimized Cap 1 mRNA can be efficiently translated into prefusion-stabilized spike protein in the primary DC (Figure 1B).

### MuVaxx delivers antigen to LNs and improves immunity against of a protein antigen

To explore the potential of delivering vaccine constituents to draining LNs, we utilized MuVaxx, our lymphatic drug delivery device, which is connectable to any luer lock syringe (Figure 2A), and which consists of microneedles that puncture the stratum corneum and release drug at the epidermal/dermal boundary. To evaluate LN delivery using MuVaxx, we used Indocyanine Green (ICG) which can be visualized *in vivo* with near-infrared fluorescence (NIRF). Following injection, we observed ICG accumulation within minutes in the draining brachial LN (Figure 2B). We then explored the potential of MuVaxx to augment the immunogenicity of a model antigen, specifically Ovalbumin (OVA) in mice. Mice were injected with OVA and an oligonucleotide adjuvant (CpG) on days 0 and 14 using an IM or MuVaxx administration. All mice treated with MuVaxx generated anti-OVA IgG by day 13 compared to 4 of 8 in the IM cohort. Additionally, following the booster shot (day 14), anti-OVA IgG was measured on day 34 and mice treated with MuVaxx displayed significantly higher titers compared to mice treated IM resulting in over a 60-fold increase in titers (Figure 2C). The cellular immune response was additionally measured in both cohorts of mice looking at cytokine production in CD8 T cells following the booster shot on days 20 and 28. Interferon-Gamma (IFNγ) and Tumor Necrosis Factor-Alpha (TNFα) were assessed following *ex vivo* stimulation with SIINFEKL, an OVA derived class I peptide. Mice treated with MuVaxx displayed higher proportions of cytokine producing CD8 T cells compared to naïve mice on both days 20 and 28 (Figure 2D). Taken together, these results highlight the potential of MuVaxx to deliver cargo to draining LNs to improve immunity of vaccines.

**Figure 2).**
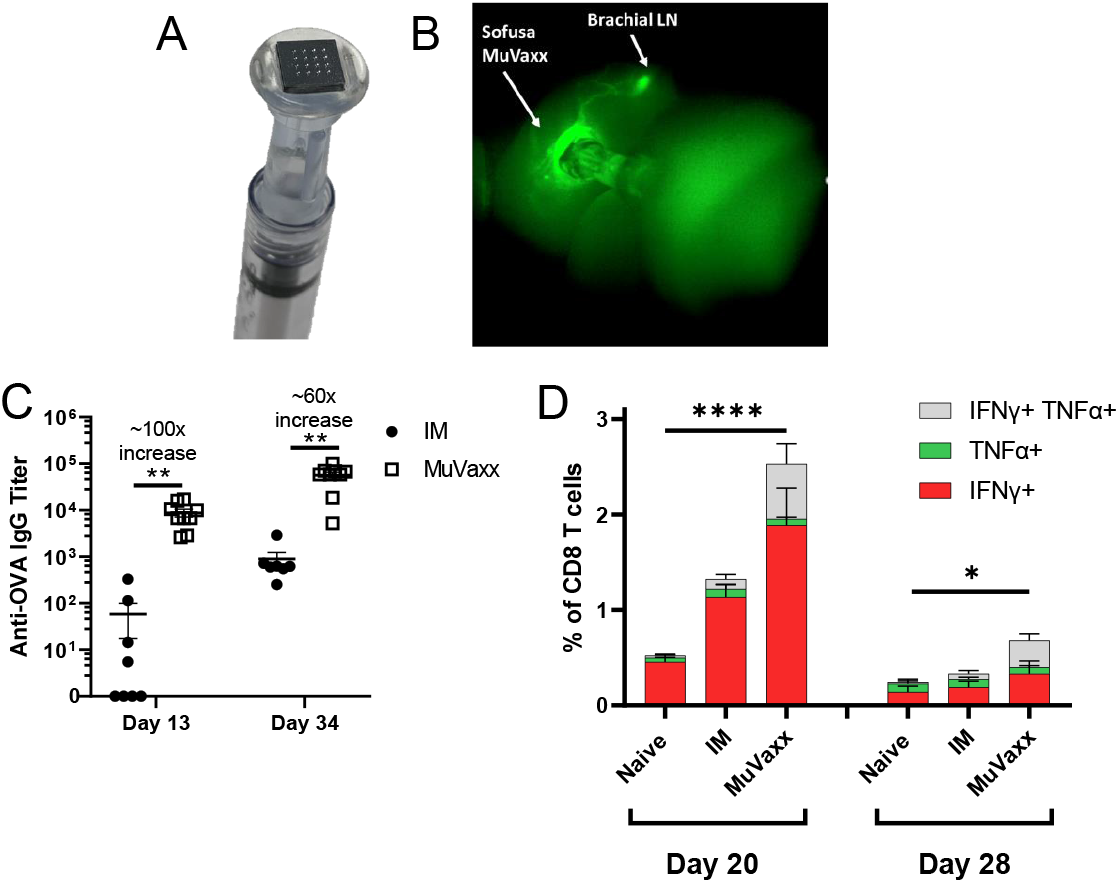
MuVaxx enables delivery to draining LNs and improves immunity compared to IM administration. A) Image of MuVaxx device connected to 1 mL syringe. B) Image of C57Bl6 mouse 5 minutes after injection of ICG using MuVaxx. C-D) C57Bl6 mice were vaccinated with 10 μg of OVA and 8 μg of CpG on days 0 and 14. C) Serum was collected on days 13 and 34 and anti-OVA IgG was quantified via serial dilutions run by ELISA assays. D) Whole blood was collected and stimulated with SIINFEKL peptide followed by ICS to measure IFNγ and TNFα in the CD8 T cell compartment. C-D) Data represent n=8-9 mice per group. Cytokine statistics represents difference between IFNγ+ groups for panel D.

### STI-7264 induces anti-spike antibodies with MuVaxx enabling dose sparing

To evaluate the potential of MuVaxx to enhance immunity, we compared the humoral immunity of STI-7264 against the Reference vaccine along with the comparison of IM vs MuVaxx administrations. C57Bl6 were immunized with either 10 μg Reference IM, or STI-7264 at 10 μg IM, 1 μg IM, or 1 μg MuVaxx on days 0 and day 35 with serum collection every 7 days (Figure 3A). Seven days after the original primer shot, both 10 μg mRNA-LNP formulations (Reference and STI-7264) administered IM showed high anti-RBD specific IgM responses with a reduction in IgM observed for the 1 μg dose. Interestingly, when 1 μg of the STI-7264 was administered using MuVaxx, a similar IgM response was measured (Figure 3B). Similar trends were also observed when quantifying the IgG response for both anti-RBD and anti-S1 specific IgG antibodies (Figure 3C-D) highlighting the dose sparing potential that may be induced with MuVaxx. Additionally, when exploring the IgG response following the initial dose, MuVaxx elicited sustained anti-Spike IgG antibodies in the serum compared to the traditional IM administration which waned at a faster rate (Figure 3E-F). Overall, these results highlight successful generation of a spike encoding mRNA vaccine with delivery toward draining LNs and/or cells within the epidermis via MuVaxx enabling dose sparing activity.

**Figure 3).**
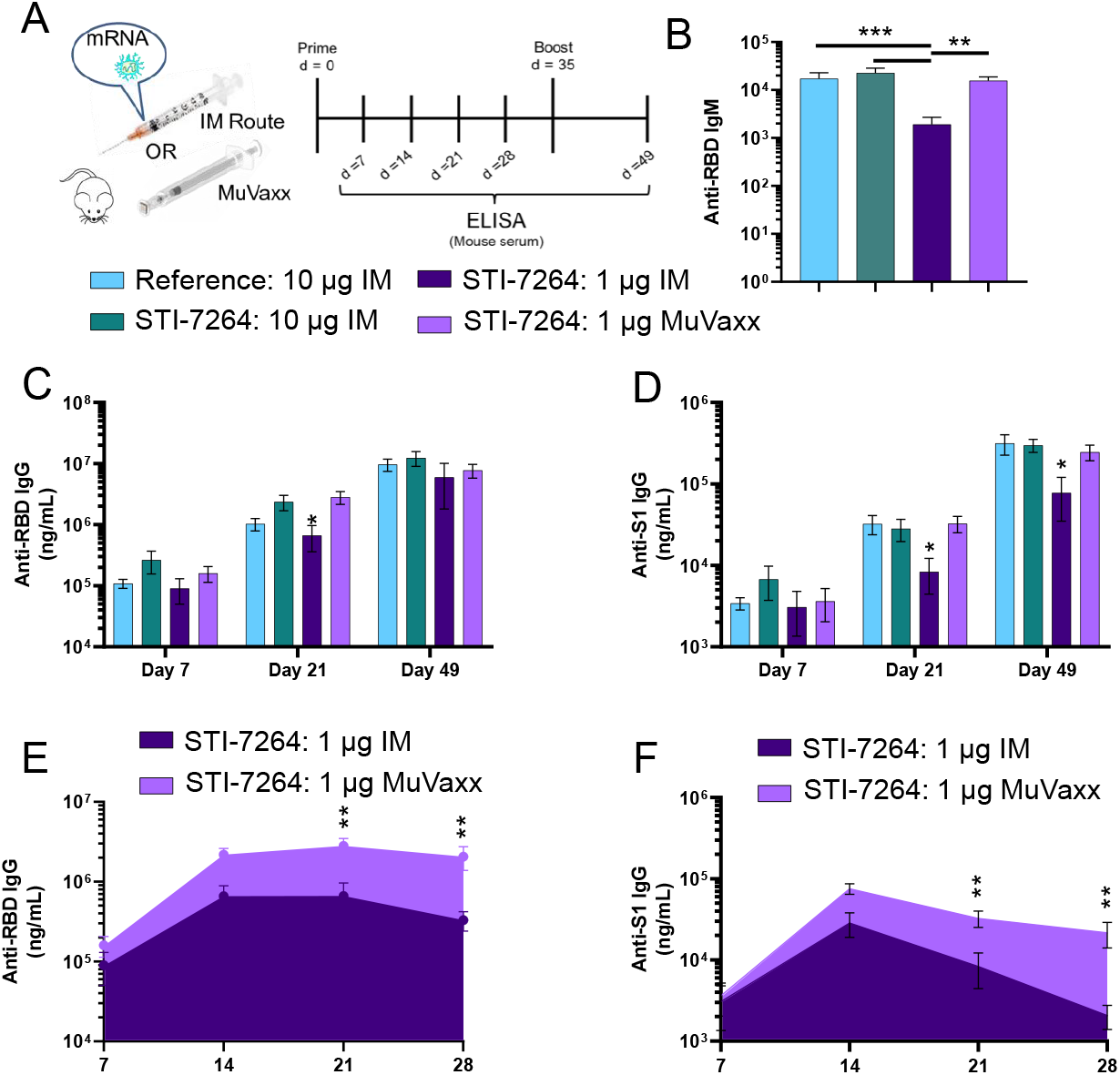
MuVaxx improves humoral immunity of spike encoding mRNA – LNP formulation. A) Treatment schedule. B-F) Serum was collected and tested for spike specific antibodies. B) Day 7 anti-RBD IgM. C) Anti-RBD IgG on days 7, 21, and 49. D) Anti-S1 IgG on days 7, 21, and 49. E) Anti-RBD IgG AUC plots comparing IM and MuVaxx 1 μg dose. F) Anti-S1 IgG AUC plots comparing IM and MuVaxx 1 μg dose. Data represent n=5 mice per group. Stats in C and D represent significance against STI 1 μg IM vs all other groups at that day.

### STI-7264 favors Th1 response over Th2

To assess the Th1/Th2 bias elicited after immunization, whole blood was collected to measure cytokine producing CD4 T cells 6 days after the booster shot via intracellular cytokine staining (ICS) and IgG subclass titers from day 49 serum (Figure 4A). The ratio of Th1 (CD3^+^CD4^+^IFNγ^+^) to Th2 (CD3^+^CD4^+^IL4^+^) T cells favored a Th1 response and was similar between all cohorts, although the 10 μg STI-7264 IM group and 1 μg STI-7264 MuVaxx groups had slightly higher ratios favoring an enhanced anti-viral immune response (Figure 4B). Similarly, the ratio of IgG2c to IgG1 was skewed towards IgG2c for the 10 μg STI-7264 IM group and 1 μg STI-7264 MuVaxx cohorts suggesting bias towards a Th1 response (Figure 4C) in line with the CD4 T cell cytokine phenotypes.

**Figure 4).**
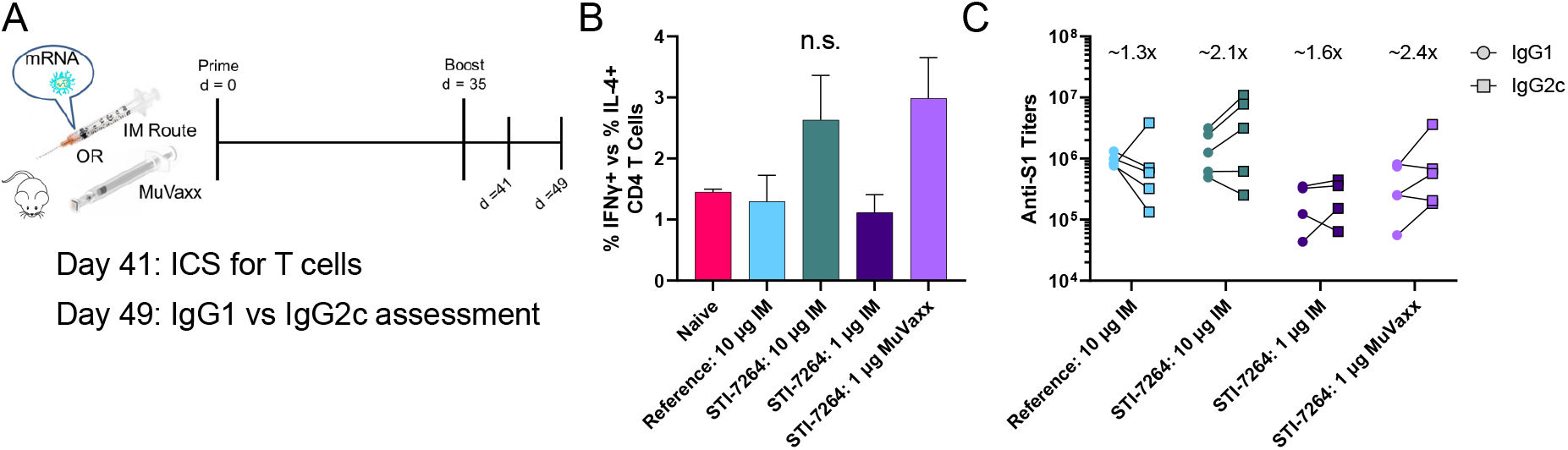
STI-7264 and MuVaxx bias response towards Th1 population. A) Treatment schedule. B) Whole blood was incubated with spike associated peptides (Miltenyi Biotec Peptivator) overnight followed by ICS. Ratio of CD4 Th1 (CD4 IFNγ+) to Th2 (CD4 IL-4+) phenotypes from whole blood on day 41 is shown. C) Serum was assessed by ELISA for S1 specific IgG1 and IgG2c on day 49 by end-point titers. Numbers listed above each group represent average IgG2c : IgG1 ratio. Data represent n=5 mice per group.

### Elevated CD8 T cell immunity is elicited following vaccination with STI-7264 and MuVaxx

In addition to CD4 T cells, the responses in the CD8 T cell compartment were evaluated on day 49 following incubation with spike associated peptides overnight via ICS (Figure 5A). Mice vaccinated with the Reference vaccine displayed a minor increase in cytokine producing CD8 T cells (Figure 5B and C), in line with previous literature^16^. However, the STI-7264 10 μg formulation when administered IM led to a robust antigen specific CD8 T cell response as measured by IFNγ and TNFα. The response was dose dependent as IM administration of 1 μg of STI-7264 led to a minimal CD8 T cell response. Interestingly, when 1 μg of STI-7264 was administered via MuVaxx toward draining LNs, the CD8 T cell response was restored to similar levels to that of a 10 μg IM dose (Figure 5B and C). Overall, these results highlight the improved CD8 T cell response observed with this STI-7264 formulation along with the benefit of directing spike encoding mRNA towards draining LNs to generate CD8 T cell immunity.

**Figure 5).**
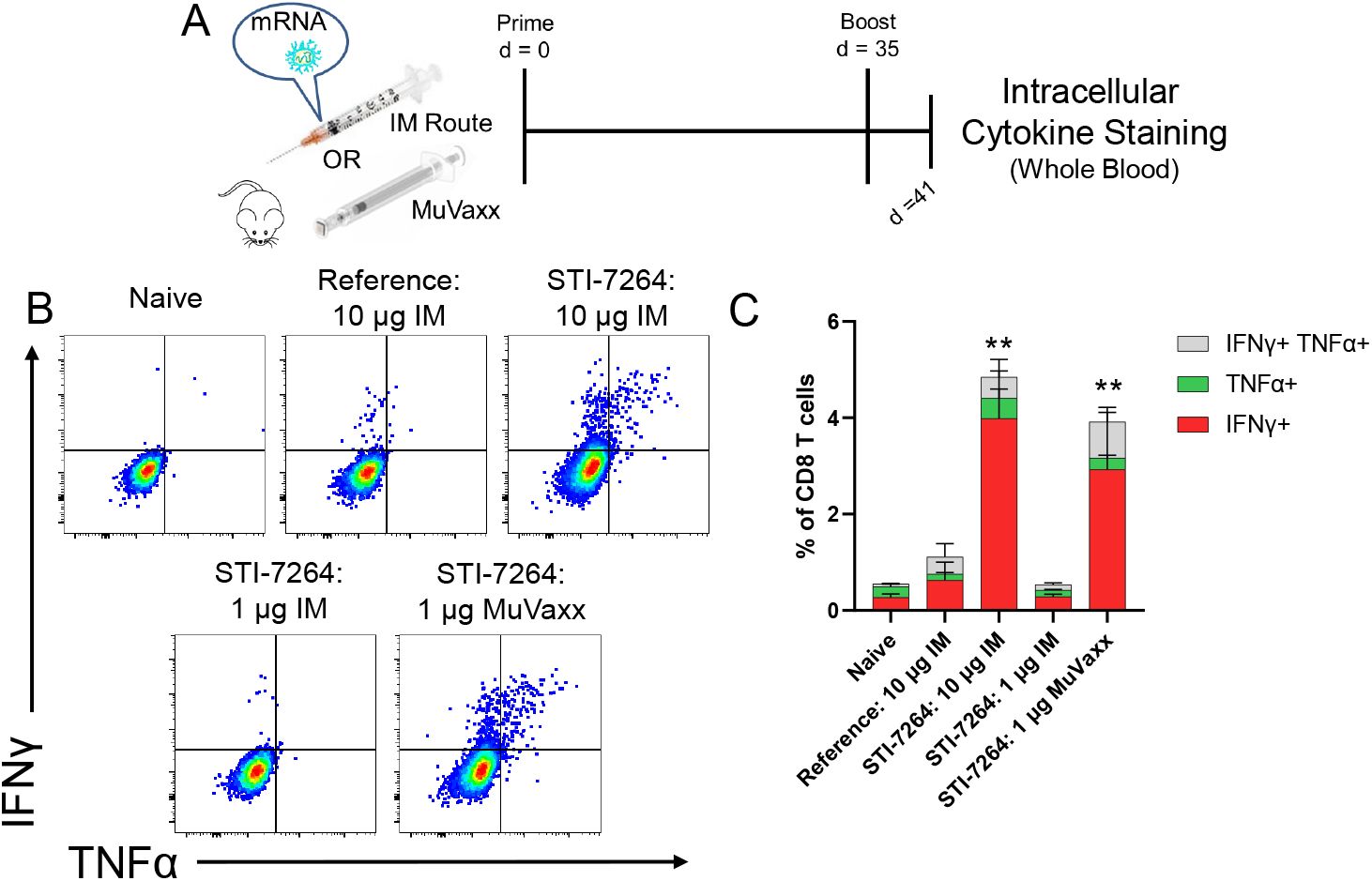
STI mRNA formulation improves CD8 T cell immunity. A) Vaccine experiment timeline: 6 days following booster shot, intracellular cytokine staining was performed in the presence of spike associated peptides (Miltenyi Biotec Peptivator). B) Representative flow cytometry plots of IFNγ and TNFα production from CD8 T cells. C) Quantification of B. Statistical analysis was done using ANOVA with Tukey’s test. **P < 0.01 and represents difference between other IFNγ+ groups. Data represent n=5 mice per group.

### Neutralizing antibodies are generated following STI-7264 vaccination

To asses nAb generation, a Plaque Reduction Neutralization Test (PRNT) was performed *in vitro*, where VeroE6 cells were exposed to the live virus in the absence or presence of diluted mouse serum. PRNT detects plaque formation and is indication of cell infection by the SARS-CoV-2 virus whereas the absence of plaque formation represents nAb presence. Each cohort of treatments led to nAb generation by d49 against the WT strain (Figure 6A). To investigate protection against the Beta variant, VeroE6 cells were incubated with this strain of the virus. The Reference vaccine, STI-7264 10 μg IM, and STI-7264 1 μg MuVaxx cohorts displayed robust protection against this strain up to a 1:120 dilution whereas the lower 1 μg STI-7264 IM cohort of mice displayed much lower protection (Figure 6B). Taken together, these results show that STI-7264 generates nAbs and that lymphatic mediated delivery via MuVaxx can broaden protection at 1/10^th^ the IM dose.

**Figure 6).**
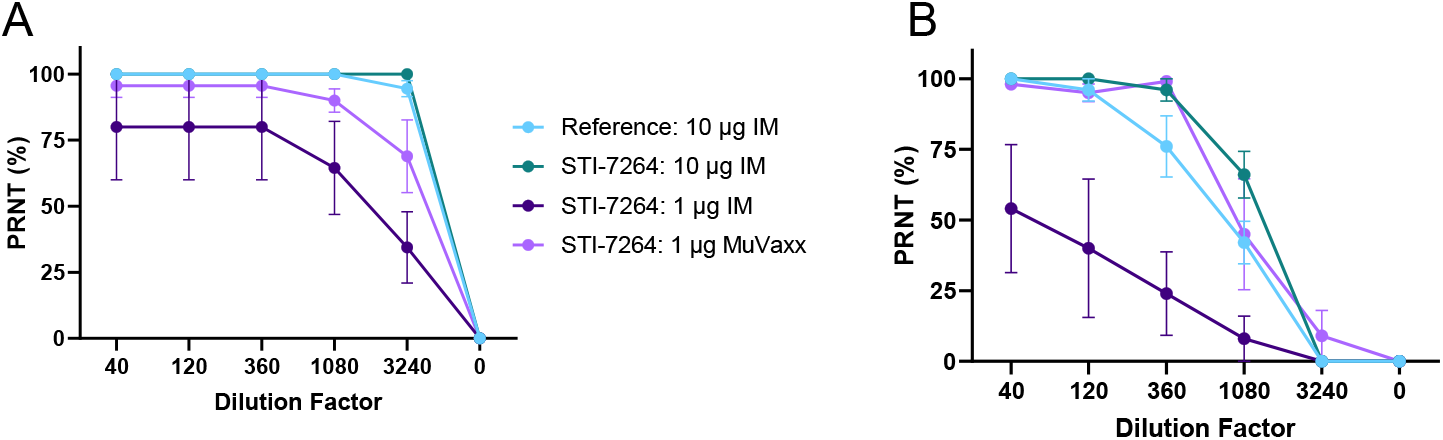
STI-7264 elicits nAbs against WT virus and Beta variant. A-B) PRNT dilution curves from mouse serum on day 49 post prime. A) PRNT curves against WT virus. B) PRNT curves against Beta variant. Data represent n=5 mice per group.

## Discussion

Here, we report a novel mRNA-based SARS-CoV-2 vaccine, STI-7264, that induces similar humoral immunity with elevated cellular immunity compared to a Reference vaccine formulation when administered IM. Immunity generated with the STI-7264 formulation was dose dependent as immunity was reduced when going from 10 to 1 μg IM. Interestingly, when administering the same 1 μg STI-7264 formulation via MuVaxx, dose sparing effects were observed where both humoral and cellular immunity were similar to that of a 10 μg IM dose highlighting the improved immunogenicity when directing vaccines towards LNs. Previous mRNA-based vaccines have reported dose-dependent side effects with higher doses linked to systemic and local adverse events^17,18^ underscoring an additional advantage of lower dose formulations; lessening side effects while expanding vaccine access to large populations.

A vaccine that generates durable immunity is another hallmark of an effective vaccine and is a metric we investigated. The serum concentrations of both anti-S1 and anti-RBD IgG waned to a lesser degree in mice treated with MuVaxx relative to those treated with the same dose IM highlighting improved durability. This is of interest as recent reports have shown declines in SARS-CoV-2 neutralizing antibodies 2-3 months after disease onset as short-lived plasma cells stop producing nAbs.^19,20^ However, a subset of plasma B cells do differentiate into memory B cells following infection and/or vaccination leading to persistent germinal center formation within LNs where somatic hypermutation takes place.^21–23^ Thus, efficient delivery of vaccines towards LNs where memory B cells reside may improve memory B cell activation and coverage against emerging variants at lower doses in line with the nAb data shown here.

While most emphasis of the available SARS-CoV-2 vaccines has been focused on the humoral response and nAbs, an ideal vaccine should also generate robust T cell immunity to synergistically protect against infections. In the context of COVID-19, subsets of patients with preexisting SARS-CoV-2 specific T cells have demonstrated rapid viral clearance and less severe disease highlighting their important role for disease prevention.^3,4^ Additionally, T cell responses against previous *betacoronaviruses* can persist for decades^5,24^ and display cross-reactivity against other *betacoronaviruses*^25–27^ highlighting potential for long term protection and coverage against variants. In this work, we show STI-7264 delivered via MuVaxx elicits a strong CD8 T cell response towards SARS-CoV-2 peptides which may be advantageous for preventing COVID-19 and providing protection against re-infection.

The CD4 Th2 phenotype has previously been associated with vaccine-associated enhanced respiratory disease (VAERD) in those vaccinated against the measles- and respiratory syncytial-virus.^28,29^ We therefore explored the CD4 Th1 vs Th2 response and IgG2c to IgG1 antibody response in vaccinated mice. The response of all vaccinated mice favored a CD4 Th1 response in line with naïve mice. In line with the T cell response, the antibody response also skewed towards a Th1 response as measured by the IgG2c to IgG1 ratio in vaccinated mice with the STI-7264 10 μg IM and 1 μg MuVaxx cohorts displaying a stronger IgG2c to IgG1 ratio. Mice treated with STI-7264 formulations (high dose IM or low dose MuVaxx) may display enhanced anti-viral activity as mouse IgG2 subclasses have been shown to induce antibody-dependent cellular cytotoxicity.^30^ Taken together, the Th1:Th2 response shown here suggests promising activity for avoiding VAERD while promoting anti-viral activity.

Overall, we show preliminary results highlighting improved immunogenicity of a novel mRNA-LNP formulation, STI-7264 with dose sparing potential demonstrated when directing towards the epidermis and draining LN using MuVaxx. Further research is needed to determine cellular and humoral durability compared to an IM administration along with coverage against known and emerging VoCs. However, the results are promising for continuing forwards and evaluating in the clinical setting for vaccinating against SARS-CoV-2.

## Materials and Methods

### *In vitro* transcription and purification of RNA

To generate the template for RNA synthesis, the sequence of the SARS-Cov-2 Spike protein (GenBank: QHD43416.1) was codon optimized and cloned into pVAX1-based backbone which features 5′-UTR, 3′-UTR and Poly-A tail. To increase the protein stability, 2P mutations at positions 986-987 were introduced. The plasmid DNA was produced in bacteria, purified and linearized by a single-site restriction enzyme digestion. The template DNA was purified, spectrophotometrically quantified, and *in vitro* transcribed by T7 RNA polymerase (Cat: M0251, NEB) in the presence of a trinucleotide cap1 analogue, m7(3OMeG)(5′)ppp(5′)(2OMeA)pG (Cat: N-7113, TriLink), and of N1-methylpseudouridine-5’-triphosphate (Cat: N-1081, TriLink) in place of uridine-5’-triphosphate (UTP). After the reaction, DNase I (Cat: M0303, NEB) was added to remove the template DNA and the mRNA was purified by LiCl precipitation (Cat: AM9480, ThermoFisher).

### In vitro mRNA expression

Monocytes are isolated and differentiated into DCs in presence of GM-CSF (Cat: 300-03, Peprotech) and IL-4 (Cat: 200-04, Peprotech). Between day 6-day 8, cells were transfected with mRNA by the Neon^TM^ electroporation transfection system (Cat: MPK5000, ThermoFisher). 24 hours post-transfection, the cells were collected and stained with anti-Spike antibody STI-2020 in FACS buffer (DPBS+0.5%BSA) for 30 minutes on ice. Thereafter, cells were washed twice in FACS buffer and incubated with rat anti-human Fc antibody conjugated to allophycocyanin (Cat: 410712, BioLegend) for 15 minutes on ice. The cells were washed with FACS buffer and analyzed by the Attune NxT Flow Cytometer (ThermoFisher).

### SARS-CoV-2 Virus

SARS-COV-2 viruses were obtained from BEI resources (Washington strain NR-52281;; Beta Variant NR-54009) VeroE6 monolayers were infected at an MOI of 0.01 in 5ml virus infection media (DMEM + 2% FCS + 1X Pen/Strep). Tissue culture flasks were incubated at 36°C and slowly shaken every 15 minutes for a 90 minute period. Cell growth media (35mL) was added to each flask and infected cultures were incubated at 36°C/5% CO2 for 48 hours. Media was then harvested and clarified to remove large cellular debris by room temperature centrifugation at 3000 rpm.

### Animals and in vivo studies

6- to 12-week old C57Bl6 mice were purchased from the Jackson Laboratory. All protocols were approved by the Institutional Animal Care and Use Committee (IACUC). Mice were injected with the indicated administration technique under isoflurane anesthesia in the right hind flank area for IM injections and in the right dorsal area for MuVaxx injections. For imaging studies, ICG (Sigma) was dissolved at 2.5 mg/mL in deionized water and injected using MuVaxx with a NIRF camera used to collect images 5 min following injection. For OVA vaccine studies, 10 μg of OVA (Cat: VAC-POVA, Invivogen) and 8 μg of CpG (Cat: TLRL-1826-1, Invivogen) were administered to mice on days 0 and 14. Peripheral blood was collected from anaesthetized mice once/week via submandibular route. Reference mRNA-LNP vaccine is the same construct as an EUA cleared compound.

### ELISA assays

To asses spike specific antibodies, S1 (Cat: 40591-V08H Sino Biological) or RBD (Cat: 40592-V08B, Sino Biological) protein was coated on half-area high binding plates (Cat: N503 Thermo) at 1 μg/mL overnight at 4°C. Plates were washed 3 times with ELISA wash buffer (Thermo), pre-blocked with casein blocker (Cat: 37528 Thermo) for 1 hour at room temperature (RT), and washed 1 time with ELISA wash buffer. Mouse sera was diluted in casein blocker and transferred to ELISA plates for 1 hour at RT followed by 3 wash steps. Secondary antibody of horse radish peroxidase (HRP)-conjugated rabbit anti-mouse IgM (μ chain), HRP-conjugated rabbit anti-mouse IgG (Fcγ), HRP-conjugated goat anti-moues IgG1, or HRP-conjugated goat anti-mouse IgG2c was added to ELISA plates for 1 hour at RT followed by six wash steps. Plates were developed with TMB substrate solution (Cat: 34021 Thermo) for approximately 10 min at RT and stopped with 2 normal sulfuric acid. The absorbance was measured at 450 nanometers using a BioTek Cytaktion 5 plate reader. For IgG (Fcγ), a standard curve was generated using Anti-RBD PAb (Cat: 40592-MP01 Sino Biological) or Anti-S1 (Cat: MAB105405 R&D Systems) starting at 3000 ng/mL with 3 fold serial dilutions. For IgM (μ chain), IgG1, and IgG2c, serial fold dilutions were run and titers were determined using an absorbance cutoff of 0.7 OD.

### Plaque Reduction Neutralization Test (PRNT)

Simian VeroE6 cells were plated at 18×10^3^ cells/well in a flat bottom 96-well plate in a volume of 200 μl/well. After 24 hours, a serial dilution of seropositive blood serum is prepared in a 100 μl/well at twice the final concentration desired and live virus was added at 1,000 PFU/100μl of SARS-CoV-2 and subsequently incubated for 1 hour at 37°C in a total volume of 200 μl/well. Cell culture media was removed from cells and sera/virus premix was added to VeroE6 cells at 100 μl/well and incubated for 1 hour at 37°C. After incubation, 100 μl of “overlay” (1:1 of 2% methylcellulose (Sigma) and culture media) is added to each well and incubation commenced for 3 days at 37°C. Plaque staining using Crystal Violet (Sigma) was performed upon 30 min of fixing the cells with 4% paraformaldehyde (Sigma) diluted in PBS. Plaques were assessed using a light microscope (Keyence).

### Peripheral blood T cell intracellular cytokine staining

ICS was performed at the indicated time points following the booster shots for IFNγ, TNFα, and IL-4. Whole blood was stimulated for 6 hours with 1 μg/mL of SIINFEKL (Sigma) or overnight with 1 μg/peptide per well of spike associated (Cat: 130-127-951, Miltenyi Biotec Peptivator) peptides at 37°C, 5% CO_2_ in the presence of brefeldin A (Biolegend) and monensin (Biolegend). Following stimulation, whole blood was incubated with red lysis buffer (Cat: A10492-01 Gibco) at room temperature. Cells were permeabilized using Intracellular Staining Perm Was Buffer (Cat: 421002 Biolegend). Cells were stained with PE anti-mouse IFNγ (Cat: 505808 Biolegend), FITC anti-mouse TNFα (Cat: 506304 Biolegend), BV421 anti-mouse IL-4 (Cat: 504127 Biolegend), APC-Cy7 anti-mouse CD3 (Cat: 100222 Biolegend), PE-Cy7 anti-mouse CD4 (Cat: 25-0041-82 Invitrogen), and allophycocyanin anti-mouse CD8a (Cat: 100712 Biolegend). Naïve mice (non-vaccinated mice) were used as negative controls. Cells were then run on a Beckman Coulter CytoFLEX instrument and analyzed via FlowJo V10 software.

### Statistics

Statistical significance of differences between experimental groups was determined with Prism software (Graphpad). All data are expressed as standard error mean (SEM). ****P < 0.0001, ***P < 0.001, **P < 0.01, and *P < 0.05 by unpaired two-tailed *t* tests or one- or two- way analysis of variance (ANOVA).

## Author Contributions

D.F., R.C., N.S., and H.X. conceptualized and designed experiments. D.F., S.K., Q.H., X.L., R.Q., W.L., H.W., H.S., N.S. H.X. performed experiments. D.F., R.C., H.X., Q.H., X.L. B.C., R.R., and H.J. analyzed and interpreted the data. D.F., R.C., and N.S. wrote the paper.

